# Reveal Principles of Codon Optimization via Machine Learning

**DOI:** 10.64898/2026.04.16.718958

**Authors:** Deng Feng, Li Hui, Sun Dashan, Duan Guangyou, Sun Zhifei, Xue Gaoxu

## Abstract

High level of protein expression is usually welcomed in industry and research, and codon optimization is widely used to achieve high expression. Methods of implementing codon optimization can be divided into two branches, one is classical methods which develop cost functions based on empirical law, another is AI methods which learn the codon choice principles from endogenous genes with neural networks. Here we develop two codon optimization tools based on two branches respectively, namely OptimWiz 2.1 and OptimWiz 3.0. Results of fusion protein fluorescence detection indicate that both OptimWiz 2.1 and OptimWiz 3.0 are superior to all the other commercially available codon optimization tools. Principles of codon optimization are revealed in the process of machine learning on both tools.

## Introduction

In the process of translation, codon functions as the basic unit of correspondence between nucleic acids and protein. Synonymous codons are codons that encode the same amino acid. Early studies reveal that altering synonymous codon usage affects protein expression level [1–2]. Based on this recognition, Sharp et al. introduced codon adaptation index (CAI) and proved that CAI values are parallel with gene expression level in many cases [3], which can be considered as the first codon optimization strategy.

Codon optimization tools developed in recent years are much more complex than just calculating CAI values. Multiple factors besides CAI and various algorithms are used in codon optimization implementation. Generally, codon optimization implementation can be divided into classical and AI methods. Classical methods are based on empirical knowledge on rules between protein expression level and synonymous codon usage. In this implementation strategy, cost functions describing empirical rules will be developed and then synonymous codon usage will be altered to reach pareto-optimal solutions. Two key-points to success in this strategy are the ability to reach pareto-optimal solutions and the accuracy that cost function describes protein expression principles. With more terms included in cost functions besides CAI, pioneers realized that codon optimization is a multi-object problem, and genetic algorithms are applied to solve this problem [4–5]. In recent years, algorithms applied to codon optimization tend to be various. Şen et al. applied mixed integer linear programming (MILP) to codon optimization in 2020 [6]. In the same year, Taneda et al. invented COSMO and displayed that COSMO is superior to genetic algorithms in finding pareto-optimal solutions [7]. In 2022, LinearDesign is invented by applying lattice parsing to mRNA design [8]. In the meantime, several variants of genetic algorithms are developed to enhance the ability of finding pareto-optimal solutions [9–11]. In this paper, we selected genetic algorithm for our codon optimization tool OptimWiz 2.1. In attempts to describe protein expression principles, factors such as CAI, context, CpG, GC ratio, MFE, CPB, HSC can be introduced to cost functions [5, 7, 12–14]. A common defect in existing methods and tools is that factors in cost functions are selected and weighted based on untested heuristics. Here we introduce a more reasonable method of cost function determination by data fitting with translatome data. Results of data fitting and wet lab validation indicate that widely accepted empirical knowledge on the role of MFE is dubious.

AI methods are based on the opinion that endogenous genes know how to choose codons. To learn from endogenous genes, neural networks are constructed and trained. In the training process, inputs are amino acid sequences and outputs are DNA sequences. Taking natural DNA sequences as ground truth, accuracy of neural networks is calculated by matching natural DNA sequences and outputs. Test set accuracy and wet lab validation are usually applied to evaluate the trained neural networks except the work by Jain et al [15]. In their work, empirical terms from classical methods such as CAI and GC content are used to evaluate the trained neural networks. It is noteworthy that most publications choose language models for codon optimization implementation. Goulet DR et al. [16] constructed a RNN model with 144 layers in total. Fu H et al. [17] constructed a 4-layer Bi-LSTM model and wet lab validations indicate that their method is competitive among commercially available codon optimization tools. Work by Gong H et al. [18] is featured by 5’UTR and 3’UTR optimization with bidirectional auto-regressive transformer. In this paper, OptimWiz 3.0 is also based on language models. During the exploration of increasing accuracy of OptimWiz 3.0, we find that genes of high expression do follow certain principles.

No matter classical methods or AI methods are selected, dataset is required for data fitting or AI training. The translatome, which includes the complete set of mRNAs actively being translated into proteins in a cell, provides a potentially valuable dataset for these purposes [19]. By focusing on the actively translated mRNAs, the translatome offers a more precise picture of the cellular protein synthesis process compared to the broader transcriptome. Techniques such as ribosome profiling have enabled detailed analysis of the translatome, uncovering translation rates and patterns of ribosome occupancy [20–21]. Utilizing translatome data might reveal specific codon usage patterns and translational efficiency, which could be beneficial for optimizing gene expression. Furthermore, translatome data can provide insights into the dynamics of translation under various cellular conditions, adding a layer of complexity and relevance to the codon optimization process. Although translatome data is affected by multiple factors, it is proved in this paper that translatome data is actually suitable for studying single factor issues.

## Material and methods

### Translatome dataset preparation

Processed ribosome footprint data and mRNA data of E.coli are downloaded from Translatome Database (www.translatomedb.net). Coding sequences (CDS) are obtained from National Center for Biotechnology Information (NCBI) with RefSeqID. Protein expression preference is evaluated by the ratio of ribosome-protected mRNA rpkm divided by mRNA rpkm.

### Model construction and training

Construction of OptimWiz 2.1 is based on genetic algorithm. A linear cost function is made up of factors including CAI, context, MFE and other elements. Weight of different factors are determined with translatome dataset by backward propagation.

Neural networks in OptimWiz 3.0 are constructed using PyTorch framework and trained with translatome dataset.

### Protein expression level quantification

GFP fusion proteins are designed for benchmark data construction and protein expression level validation. Proteins from diverse species with different functions and locations are selected to fuse with GFP. Protein linker screen pipeline is developed in PyRosetta. Protein solubility and stability is optimized using pipelines in PyRosetta [22] and ProteinMPNN [23].

CDS of different GFP fusion proteins are cloned into Pet_24a backbone. Plasmids are transformed into E.coli BL21 competent cells. After transformation, colonies are inoculated into LB medium and incubated for 16 hours. To enable the same initial condition for all samples, a 1:100 inoculation from medium is performed, cells are incubated for another 16 hours before fluorescence detection. IPTG is not added, protein of interest is expressed by leakage only. Green fluorescence detection is performed with 485/20 nm excitation and 528/20 emission.

## Results

### Design of GFP fusion proteins

Quantitative data are required for training, adjusting and evaluating models. Fluorescence intensity detection is a cheap and convenient method to quantify protein expression level. Among all kinds of GFPs, super folder [24] is selected for its outstanding solubility and stability. Previously established GFP fusion proteins [24] are used for OptimWiz 2.1 protein expression quantification before data fitting. In order to obtain more GFP fusion proteins, a pipeline of fusion protein design is developed. For each protein of interest, hydrophobic residues on the protein surface are selected using layer selector in PyRosetta and then replaced randomly with polar residues. After which, fast relax protocol in PyRosetta is applied to adjust backbone and sidechain conformation. To make up the trade-off relationship between solubility and stability, solubility optimization described above is performed several hundred times and the optimized protein with lowest energy score is selected. We also try ProteinMPNN for protein sequence optimization following a given simple monomer design pipeline. Proteins of interest are later fused to N-terminal of GFP with a linker. A protein linker screen pipeline similar to the work by Kuhlman et al. [25] is developed, commonly used linkers [26] are screened and the most suitable candidate is determined. Fluorescence intensity of GFP fusion proteins designed by our pipeline is illustrated as Table 1. We find that 17 of 20 designed GFP fusion proteins are good enough for benchmark data construction.

**Table 1.**
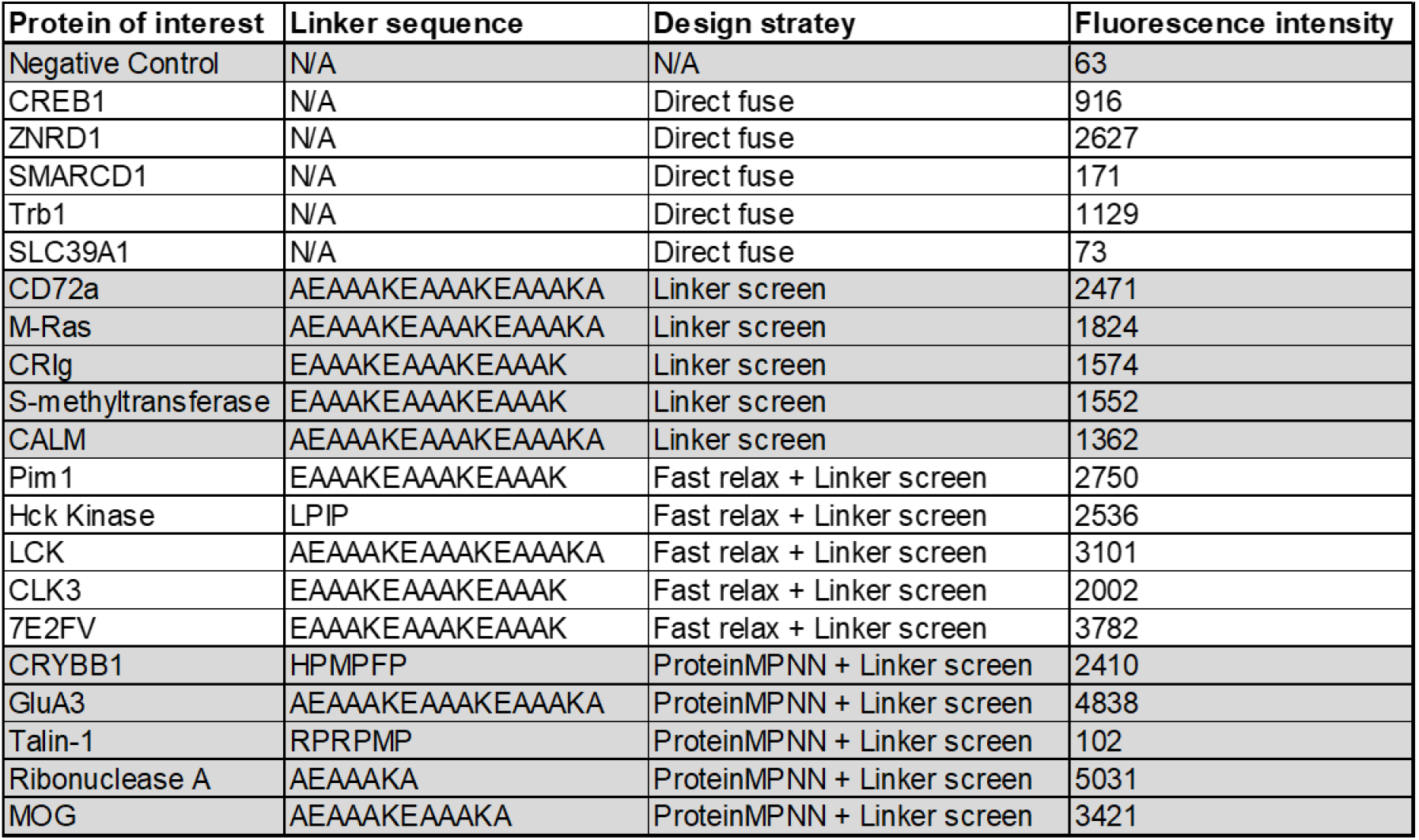
Fluorescence intensity of designed GFP fusion proteins.

### Development of OptimWiz 2.1

For codon optimization methods based on empirical knowledge, a cost function containing multiple terms is used to describe the rule of protein expression. In OptimWiz 2.1, we developed a cost function made up of CAI, context, MFE, GC ratio, Splicing Site, CpG island and other terms (equation 1). The optimized sequence is expected to satisfy all terms in cost function while these terms usually conflict with each other. To achieve the optimal solution, genetic algorithms can be applied.

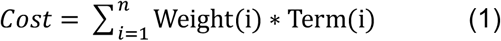

Genetic algorithms start from randomly setting a parent population (P0) and then recurrently perform recombination, mutation and sorting until the cost cannot be further decreased (Fig. 1A). In this process, recombination is the main force for pushing solutions to pareto-optimal solutions and recombination only makes sense when population diversity exists. Early versions of genetic algorithms usually lose population diversity in very limited rounds of sorting, thereby the optimization will be stopped at a point far from ideal. Advanced genetic algorithms are developed to solve this problem. In advanced genetic algorithms (Fig. 1), population diversity can be maintained by recognizing and reserving elite individuals associated with reference points.

**Figure 1.**
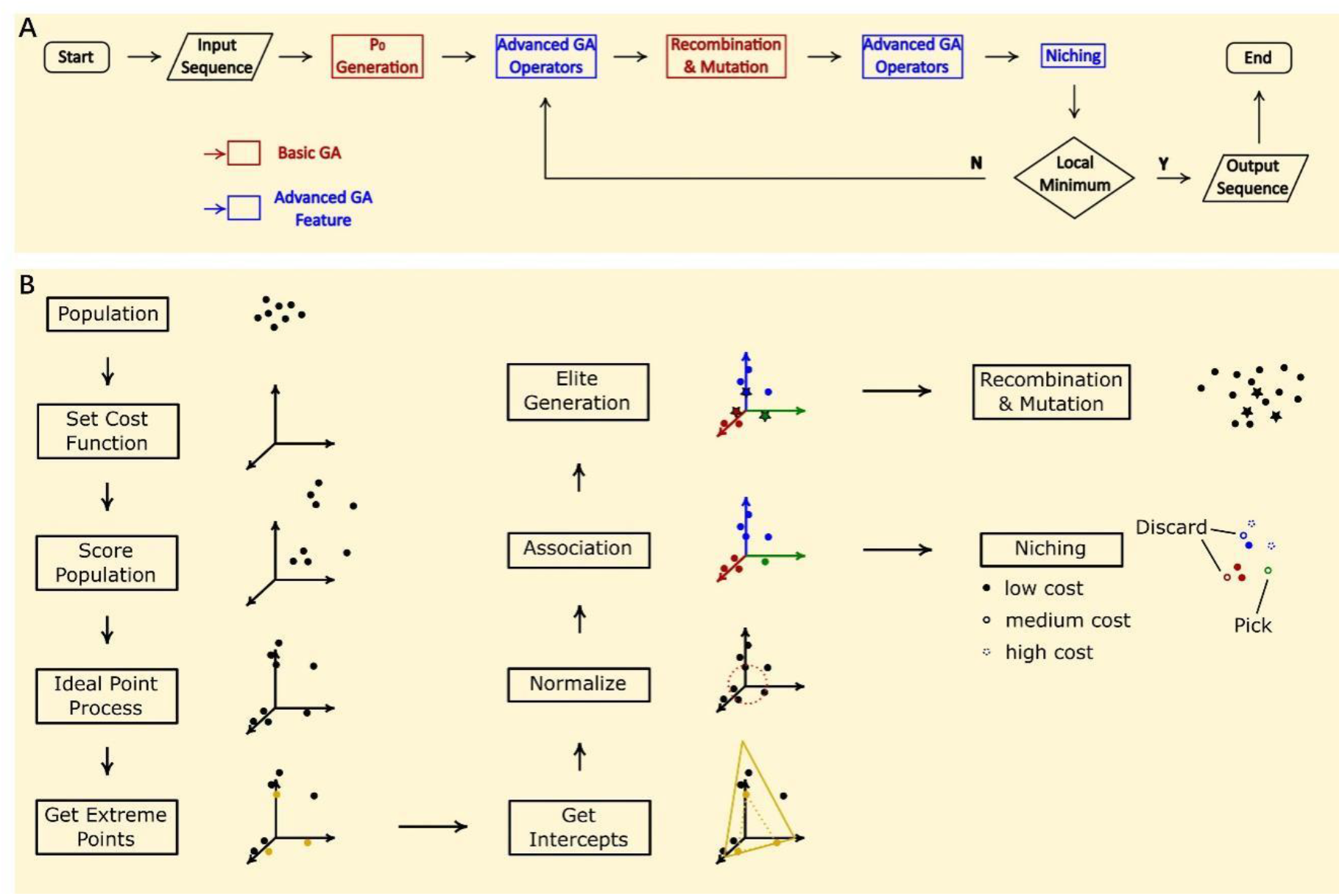
Flow chart for advanced genetic algorithm. (**A**) Flow chart for advanced genetic algorithm in general. Advanced genetic algorithm is developed based on the basic genetic algorithm. Feature of basic genetic algorithm is colored in red, feature of advanced genetic algorithm is colored in blue. (**B**) Detailed flow chart for advanced genetic algorithm feature. Advanced genetic algorithm maintains population diversity during iterative process by recognizing and reserving elite individuals.

Reaching pareto-optimal solutions of cost function is important but still not enough for success. Another key point is that cost function must properly describe the rules of protein expression. Although cost function is based on empirical knowledge, term choice and weight do not necessarily have to be decided by untested heuristics. Our strategy is to make the cost function learn the protein expression rule from benchmark data by data fitting. The benchmark data is made up of fluorescence intensity of GFP fusion proteins illustrated above, and we will later prove that translatome data is also good for use. For the first round of benchmark data construction, 5 different proteins are fused to GFP respectively. For each GFP fusion protein, commercially available codon optimization tools and OptimWiz 2.1 before data fitting are applied to generate coding sequences. Fluorescence intensity for each protein group is illustrated as Fig. 2. Equation 2 illustrated below is simple and straightforward for evaluating different codon optimization tools. Bias for third-party company / other tool 1 to 5 are 0.584, 0.230, 0.979, 0.487, 0.384 respectively. Performance of OptimWiz 2.1 before data fitting is at intermediate level with a bias equal to 0.553.

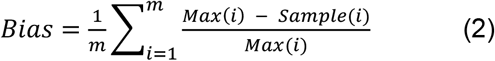

Where m is the total number of protein groups, Max refers to the maximum fluorescence intensity in this protein group, Sample refers to the fluorescence intensity of the sample being observed.

**Figure 2.**
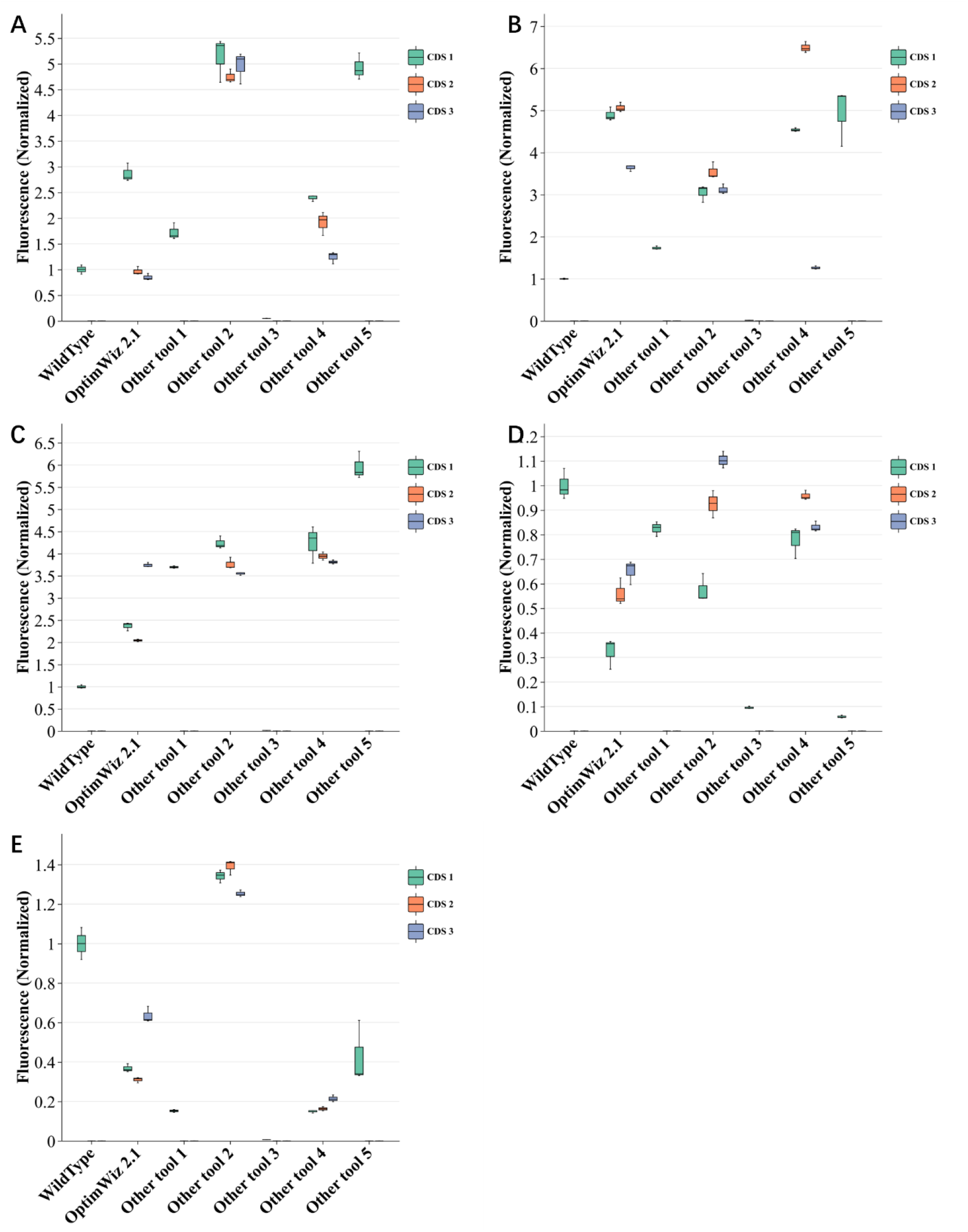
Fluorescence intensity for each protein group with different coding sequences from wild type and commercially available codon optimization tools. Data fitting is not performed yet for the OptimWiz 2.1 used here. (**A**) sulfite reductase subunit beta - GFP. (**B**) IF-5A - GFP. (**C**) chorismate mutase - GFP. (**D**) KRas4b - GFP. (**E**) TAP1 - GFP.

To improve the performance of OptimWiz 2.1, we need to first construct benchmark data. Fluorescence intensity and protein expression level are positively correlated, although this relationship is known to be not linear, fluorescence data is good enough for providing the optimization direction needed by genetic algorithms. As optimization direction in genetic algorithms is from high cost to low cost, for each protein group, cost of coding sequence with highest fluorescence intensity is assigned to 20, cost of coding sequence with lowest fluorescence intensity is assigned to 200, other coding sequences are processed linearly between the highest and lowest cost.

For each coding sequence as input, the cost function (equation 1) will generate a cost as output. Weight for each term in cost function can be adjusted to make the output as close to benchmark data as possible. Because the linear cost function is derivable, backward propagation is applied to conduct data fitting. During the process of backward propagation, negative weight is avoided by setting negative values to 0. In this way, we can put whatever we want to try into the cost function without bothering if this term really affects protein expression. Result of data fitting will tell us terms are important or not, because weight of important terms will be fitted to be high while weight of irrelevant terms will be close or equal to 0.

After data fitting, OptimWiz 2.1 is tested together with OptimWiz 3.0 and other tools on another group of GFP fusion proteins. Fluorescence intensity is given as Fig. 3. Performance of codon optimization tools is evaluated by equation 2. Bias for third-party company / other tool 1 to 5 are 0.561, 0.357, 0.866, 0.420 and 0.537 respectively. OptimWiz 2.1 after data fitting is superior to other commercially available tools with a bias equal to 0.341. OptimWiz 3.0 based on AI algorithms achieved the best overall performance with a bias equal to 0.138.

**Figure 3.**
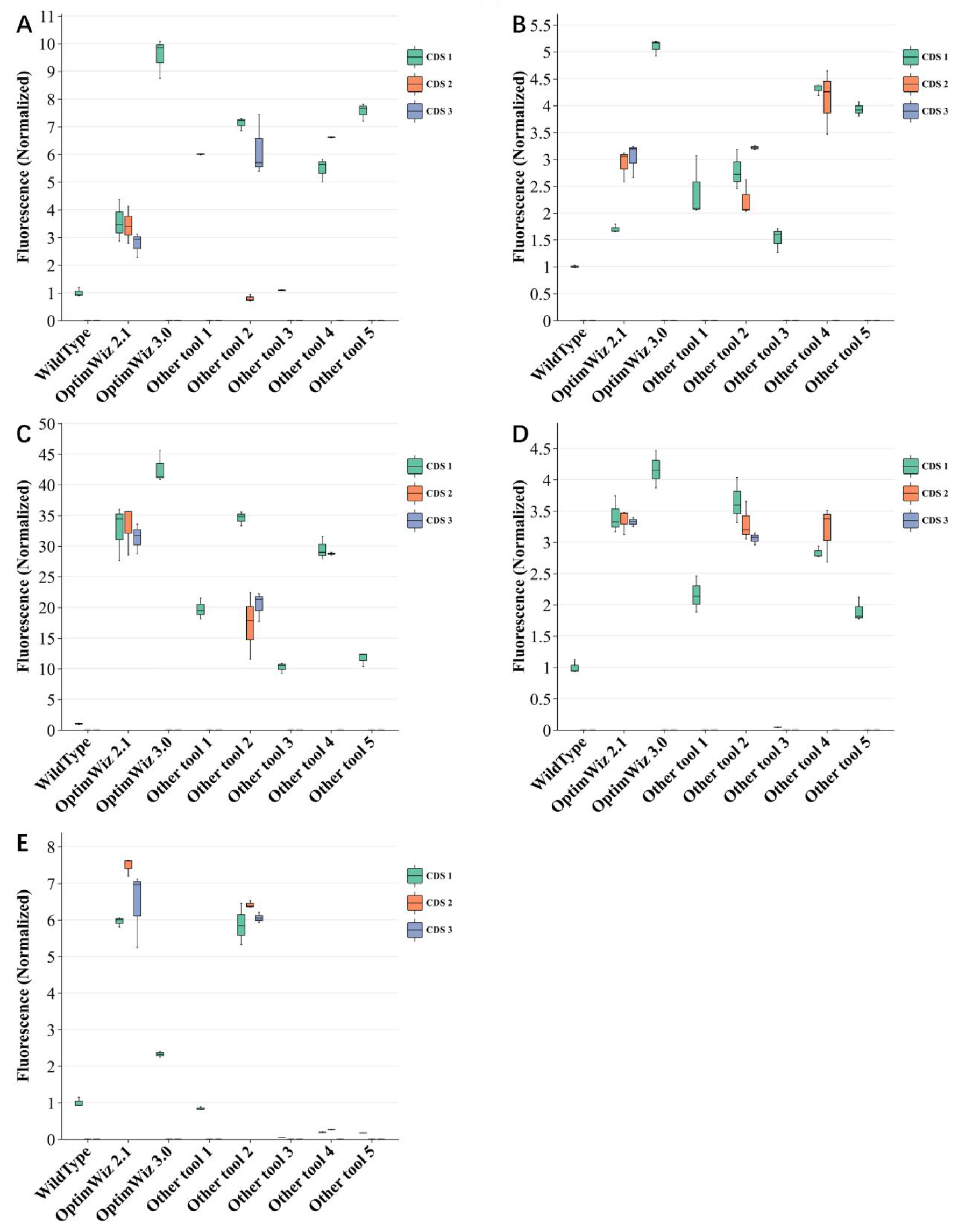
Fluorescence intensity for each protein group with different coding sequences from wild type and commercially available codon optimization tools. Data fitting is performed for OptimWiz 2.1. OptimWiz 3.0 achieved the best overall performance. (**A**) CD72 - GFP. (**B**) M-Ras - GFP. (**C**) Pim1 - GFP. (**D**) 7E2FV - GFP. (**E**) GluA3 - GFP.

### Data fitting with translatome data

Although fluorescence intensity detection is cheap and convenient compared to other protein quantification methods, gene synthesis is expensive and limits the data size. Using RNC-seq(sequencing of translating mRNA) or Ribo-seq (ribosome profiling), we are able to obtain translatome data in limited budget and estimate the expression level of thousands of endogenous proteins. Here we prove that translatome data is good for studying codon optimization, and also provide an example of exploring the role of MFE in protein translation.

Benchmark data constructed by gene synthesis is of course ideal for studying codon optimization, because the coding sequence is designed as the control variable in each protein group. In translatome data, coding sequence is not the only factor contributing to the difference in protein expression level. There are other variables like promoter, RBS, and UTR. For a cost function containing n terms (equation 1), when coding sequence is given, term values T1 to Tn and real cost negative correlated with protein expression level are fixed. Data fitting is trying to adjust the term weights to minimize deviation between estimated cost and real cost. When benchmark data is applied, the real cost is only relevant with coding sequence (equation 3). While in translatome data, other factors X also contribute to the value of real cost (equation 4). For the distribution of influence from other factors, we can make a bold assumption that it follows a normal distribution just like other natural data (equation 5). Thus, when data size m is large enough, mean value of other factors X approaches zero. By expanding Equation (4), it can be shown that the contribution of X introduces an additive constant term proportional to its variance a^2^. As a result, Equation (4) differs from Equation (3) only by a constant offset that is independent of the parameters W1 to Wn, Therefore, minimizing Equation (3) and Equation (4) yields the same optimal estimates of W1 to Wn. In another word, when data size is large enough, translatome data will be good for data fitting on a codon optimization cost function.

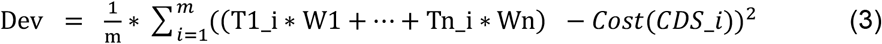

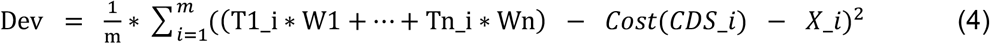

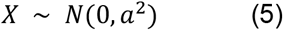

Where T1 to Tn stand for term values of a given coding sequence. W1 to Wn stand for the weights of each term. Influence from coding sequence on protein expression level is represented as Cost(CDS). Influence from other factors on protein expression level is represented as X.

We downloaded E.coli translatome and mRNA sequencing data from Translatome Database (http://www.translatomedb.net/) [27]. Coding sequences with wrong start codon, wrong stop codon or length cannot be divided by 3 are kicked out. 3637 coding sequences are obtained finally. Protein translation preference is estimated by dividing translatome rpkm with mRNA rpkm (equation 6). Protein with highest translation preference is assigned a cost of 20. Protein with lowest translation preference is assigned a cost of 200. Cost of other proteins are processed linearly between 20 and 200 by translation preference. Because we find that high expression genes follow certain principles in synonymous codon choice when training OptimWiz 3.0, which will be introduced later in this paper, 1521 sequences with preference larger than 2 are used for calculating single codon usage frequency and paired codon usage frequency. Single codon usage frequency and paired codon usage frequency will be applied for RSCU and Context calculation in cost function. After translatome data construction, data fitting is performed. Apart from adjusting the weight of each term, method of term value calculation also needs to be explored. Coefficient of determination (R^2^) is calculated to evaluate whether the adjusted cost function works better or not.

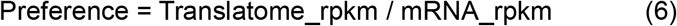

In the process of adjusting cost function, we find that the role of minimum free energy (MFE) in translation contradicts with commonly accepted point of view. It’s widely accepted that lower MFE will stabilize mRNA structure and therefore enhance protein expression [7, 28–30]. There’s an opposite view holding that rigid mRNA structure with low MFE will inhibit translation initiation and therefore decrease protein expression. To the best of our knowledge, there’s no publication on the opposite view, but some commercially available codon optimization tools are developed based on the opposite view. To figure out the role of MFE, we made three different assumptions that MFE is irrelevant with protein expression, high MFE results in low expression (high cost), high MFE results in high expression (low cost). For each assumption, MFE calculation on full length, N-terminal and C-terminal are tested respectively. Cost function is adjusted and tested based on each assumption. MFE is calculated with a method similar to the work [31] by Lorenz et al. For the assumption that high MFE gives high cost, MFE term value is calculated as equation 7. For the assumption that high MFE gives low cost, MFE term value is calculated as equation 8.

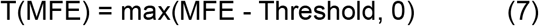

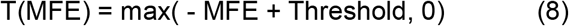

Both benchmark data and translatome data are applied to test these three assumptions. Results of data fitting on translatome data are illustrated as Fig. 4A, 4C, 4E. Results of data fitting on benchmark data are illustrated as Fig. 4B, 4D, 4F. As shown in Fig. 4, MFE calculation on the entire coding sequence fail to increase R^2^, while MFE calculation on N-terminal and C-terminal increases R^2^ only in the assumption that high MFE results in high expression (low cost). The pattern of change in R^2^ in translatome data is in line with the pattern in benchmark data. This result supports the opinion that high MFE results in high expression. Although theories on mRNA stability make sense, it seems like translation initiation has a bigger impact on protein expression level.

**Figure 4.**
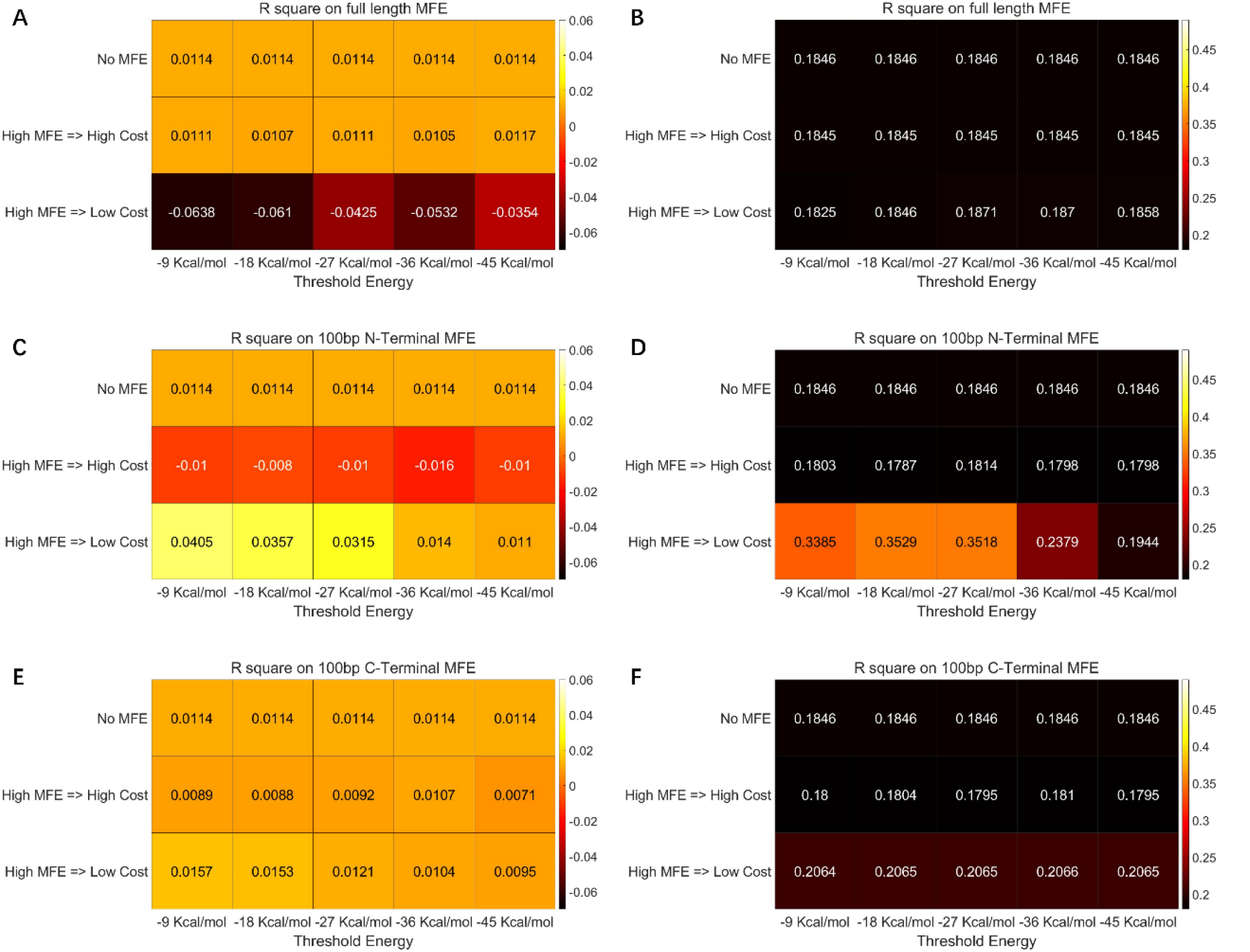
Result of data fitting on three different assumptions: MFE is irrelevant with protein expression, high MFE results in low expression (high cost), high MFE results in high expression (low cost). R^2^ is calculated to evaluate different assumptions. (**A**) MFE is calculated on full length with translatome data. (**B**) MFE is calculated on full length with benchmark data. (**C**) MFE is calculated on 100bp N-terminal with translatome data. (**D**) MFE is calculated on 100bp N-terminal with benchmark data. (**E**) MFE is calculated on 100bp C-terminal with translatome data. (**F**) MFE is calculated on 100bp C-terminal with benchmark data.

After a general conclusion on the role of MFE is drawn, we further explore the details of MFE calculation in cost function. Data fitting and R^2^ calculation are performed with different sequence length and energy threshold. Example of N-terminal MFE detail exploration is illustrated as Fig. 5. Optimum R^2^ is achieved when sequence length is set to 130bp on both translatome data and benchmark data. The most suitable threshold energy is determined as −2 Kcal/mol and −30 Kcal/mol on translatome data and benchmark data respectively.

**Figure 5.**
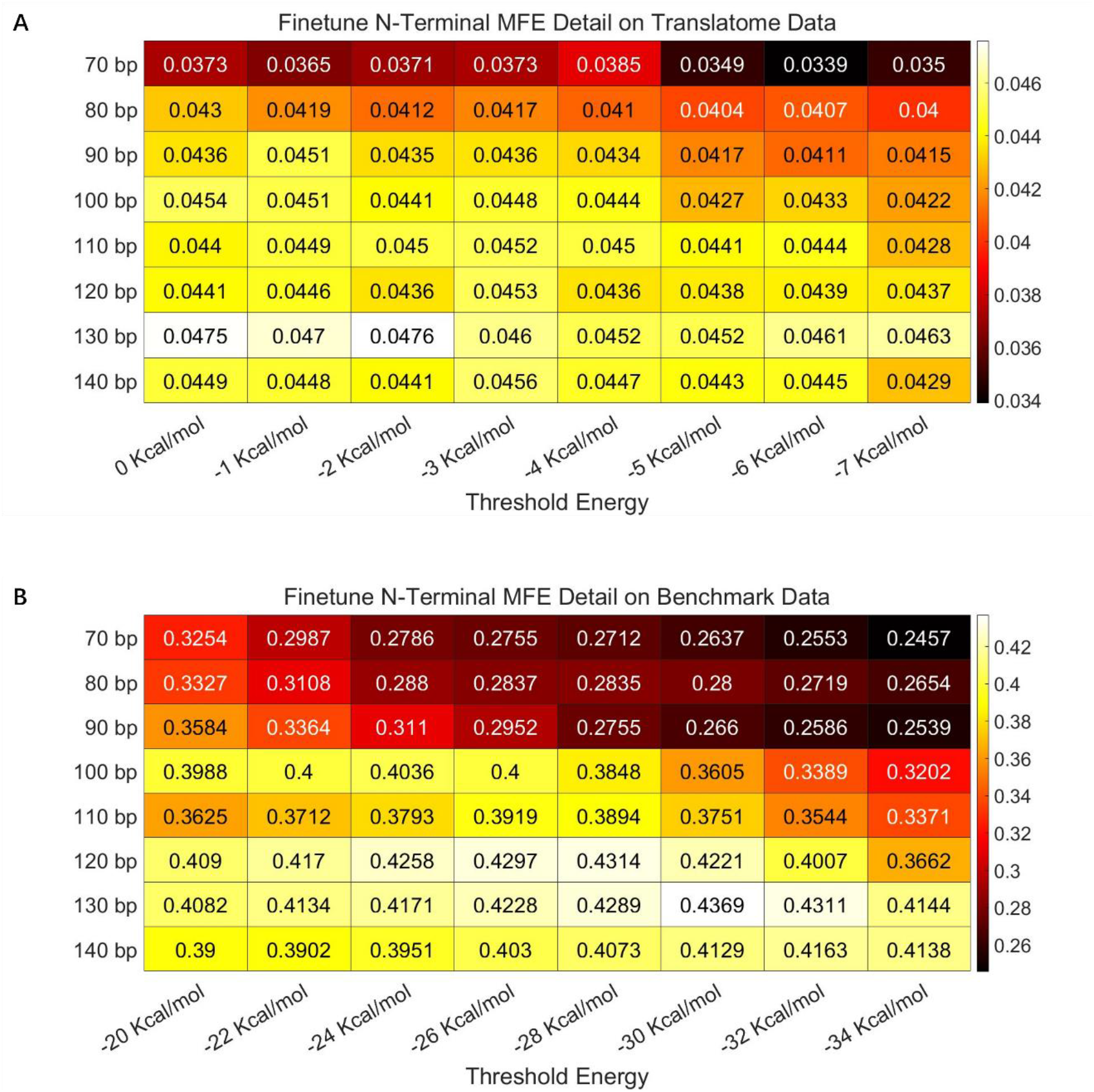
Exploration on the details of MFE calculation. (**A**) Data fitting and R^2^ calculation on translatome data. Optimum R^2^ is achieved with 130bp sequence length and −2 Kcal/mol threshold energy. (**B**) Data fitting and R2 calculation on benchmark data. Optimum R2 is achieved with 130bp sequence length and −30 Kcal/mol threshold energy.

Apart from MFE, other terms in cost function are carefully adjusted using a similar method on both translatome data and benchmark data. R^2^ on translatome data is increased from 0.0015 to 0.0571 after adjusting cost function and data fitting. R^2^ on benchmark data is increased from 0.1844 to 0.4856 after adjusting cost function and data fitting. R^2^ on translatome data is much smaller than R^2^ on benchmark data because other factors X (equation 4) would make sum of squared residuals (RSS) big. Fitted Weights for terms by translatome data and benchmark data are illustrated as Table 2. It is noteworthy that result of data fitting on term calculation detail and term weight with translatome data generally matches with the result of data fitting with benchmark data, which indicates that a throughput of several thousand data will make translatome data good enough. Considering that result of data fitting with translatome data and benchmark data still differs in some detail, and distribution of benchmark data can be biased because of its small data size, OptimWiz 2.1 finally adopted the cost function adjusted with translatome data. Result of fluorescence intensity validation (Fig. 3) indicates that adjusted OptimWiz 2.1 is superior to all other commercially available codon optimization tools.

**Table 2.** Fitted weights for terms in cost function.

|  | W(CAI) | W(Context) | W(GC) | W(MFE N) | W(MFE C) | W(Other 6) | W(Other 7) | W(Other 8) | W(Other 9) |
| --- | --- | --- | --- | --- | --- | --- | --- | --- | --- |
| Translatome data | 2.34 | 4.7 | 0.01 | 1.28 | 0.56 | 1.56 | 0.34 | 1.7 | 0.79 |
| Benchmark data | 2.25 | 3.88 | 0.06 | 0.93 | 0.53 | 2.93 | 0.36 | 3.62 | 0.73 |

Although R^2^ is increased to 0.4856 and wet lab performance gets better after data fitting, OptimWiz 2.1 is still not perfect. As we know, when a proper model is selected to describe a biological process, R^2^ can be fitted to be close to 1 [32]. Additionally, wet lab performance of OptimWiz 2.1 is not robust enough, expression level of some coding sequences optimized by OptimWiz 2.1 is only at medium level. The reason is clear that rule of protein expression is very complicated, while model with only one layer of linear cost function is far from enough to describe this rule. In order to better describe the rule of protein expression, more powerful models such as AI models are worth trying.

### Development of OptimWiz 3.0

High throughput of wet lab data supported by advanced biotechnology and development of AI algorithms, especially language models, enable AI implementations on codon optimization tasks. AI implementation on codon optimization is based on the opinion that endogenous genes know how to choose synonymous codons, and neural networks are constructed to learn from these endogenous genes. Input to neural networks are amino acid sequences and outputs are DNA sequences. Accuracy of neural networks is calculated by matching natural DNA sequences and outputs. Among publications in this field, a test set accuracy around 0.52 is usually obtained [16–17]. An accuracy of 0.63 achieved by Gong et al. [18] is commendable, but we still have to be careful that redundant sequences in hg19.knownGene dataset must be removed before training.

To develop our OptimWiz 3.0, we constructed and tested almost all kinds of neural networks such as FNN, CNN, Transformer and other language models. The dataset is made up of E.coli genome coding sequence as described by previous publication [16–18]. The finally determined network structure is featured by stacking the network unit made up of one layer of Bi-RNN and one layer of Bi-GRU. OptimWiz 3.0 consisting of two (Bi-RNN + Bi-GRU) units and two FNN layers achieved an accuracy of 0.55. Intuitionally, Bi-RNN is good at processing short term relationships such as local CAI and GC ratio, Bi-GRU is good at processing long term relationships such as avoiding splicing sites.

When genome coding sequence is used in training, we are expecting the AI model to learn from all endogenous genes. A defect in using all endogenous genes is that not every endogenous gene has high expression level. As high expression genes and low expression genes cannot be separated, while codon optimization aims to increase expression level solely, the direction that model is trained to can be somehow biased from the purpose. Since we have proved that translatome data are good for use as before, OptimWiz 3.0 is then trained with translatome data. Genes can be separated by expression level according to translation preference (equation 6). With admittable translation preference increasing from 0 to 2, test set accuracy increases from 0.55 to 0.578 (Fig. 6A). Higher admittable translation preference decreases data size and exacerbates over fitting, while test set accuracy still increases, which means that proteins of high expression do follow certain principles and these principles are learned by OptimWiz 3.0. Test set accuracy reaches 0.578 at most (Fig. 6B), further increase in translation preference decreases accuracy because of severe overfitting. Wet lab validation (Fig. 3) indicates that OptimWiz 3.0 is currently the best codon optimization tool with the bias (equation 2) equal to 0.138.

**Figure 6.**
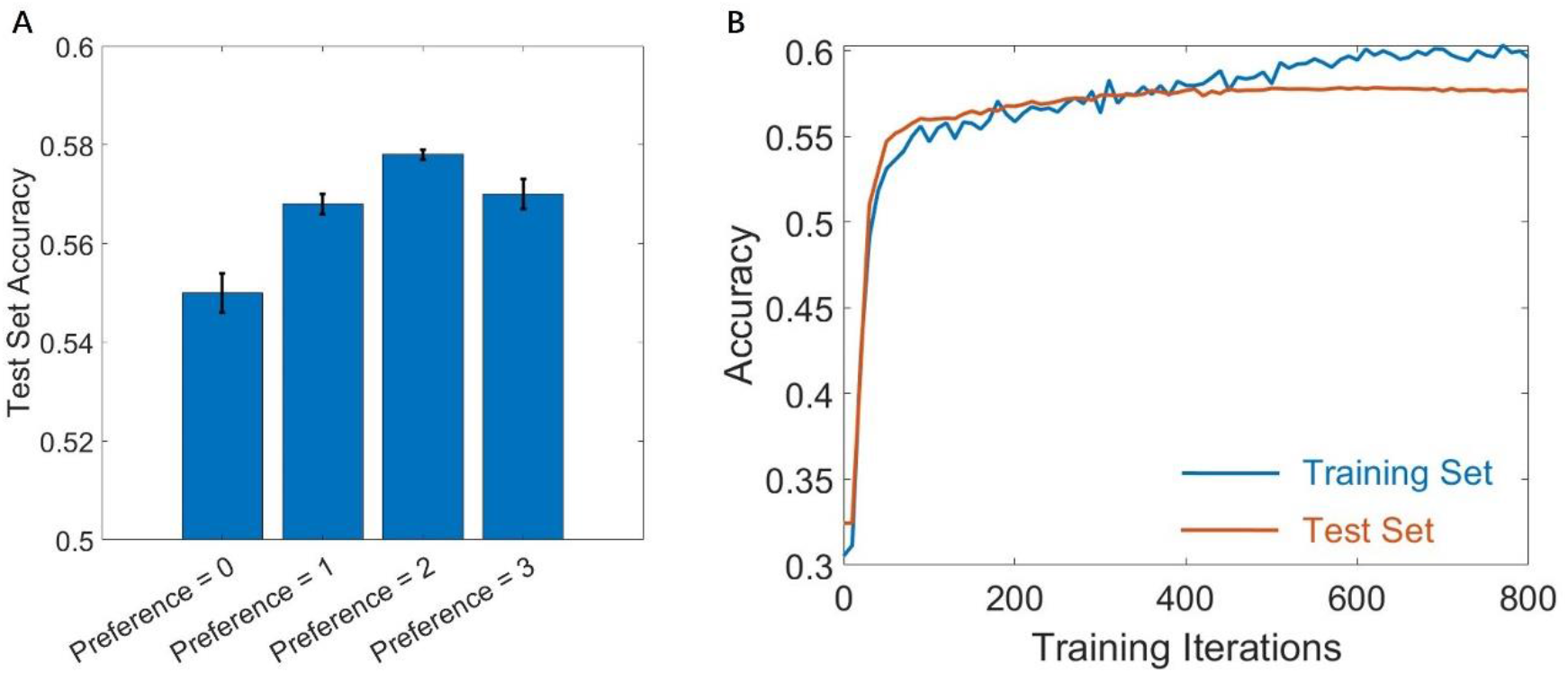
Train OptimWiz 3.0 with translatome data. (**A**) When admittable translation preference is set to 0, 1, 2 and 3, test set accuracy reaches 0.55, 0.568, 0.578 and 0.57 respectively. (**B**) Training process of OptimWiz 3.0 when admittable translation preference is set to 2. Test set accuracy reaches 0.578 at most.

## Discussion

Empirical laws are applied in various fields of biological research and manufacture. One of the reasons that empirical laws are welcomed is that they are usually intuitively easy to understand. However, empirical laws can be wrong. It is widely accepted that low MFE will stabilize mRNA and thereby yield high protein expression. In contrast, there are few if not no publications reporting that high MFE on N-terminal may make translation initiation easier. In this paper we reveal that, at least in E.coli, translation initiation is generally more important than mRNA stability in protein expression. It is beyond our expectation that high MFE on C-terminal of mRNA can help expression. To the best of our knowledge, there is currently no theory that can be applied to explain this finding. Apart from MFE, small weight of GC ratio term is another counter-intuitive finding. With these counter-intuitive findings included in the adjusted cost function, performance of OptimWiz 2.1 is better than all the other commercially available codon optimization tools. Therefore, we believe that there are still many unknown fields to be explored in biology, and empirical laws must be carefully checked before being applied to codon optimization and other tasks.

Performance of OptimWiz 3.0 is outstanding. Compared to OptimWiz 2.1 which still partially relies on empirical laws to construct cost function, OptimWiz 3.0 is a pure product of machine learning. Expression level of genes optimized by OptimWiz 3.0 is high and robust, which indicates that machine learning is more reliable than empirical laws. In previously reported AI methods of implementing codon optimization, it remains an assumption that endogenous genes know how to choose synonymous codons. It is interesting to reveal that high expression genes do follow certain principles by training OptimWiz 3.0 with translatome data. In this paper we proved that, although translatome data is affected by various factors, it is still good enough to be applied to study a single factor. It is noteworthy that this single factor is not limited to synonymous codon choice. We believe that translatome data has great potential in a wild range of applications, such as promoter, UTR and RBS design.

## Conflict of interest

OptimWiz 2.1 is protected by patent CN106951726A.

OptimWiz 3.0 is protected by patent PY24DX07369FNPE-CN.

## Author contribution

Deng Feng and Li Hui designed and performed experiments in CHO and HEK cell lines.

Sun Dashan and Xue Gaoxu designed experiments in E.Coli.

Sun Dashan designed and implemented algorithm for OptimWiz 2.1 and OptimWiz 3.0, performed experiments in E.Coli.

Duan Guangyou did code review.

Sun Zhifei leaded wet lab data collection for model training.

## Notes

### Summary of Updates

Symbols corrected in chapter 'Data fitting with translatome data'.

## References

1. Gouy M & Gautier C. Codon usage in bacteria: correlation with gene expressivity. Nucleic Acids Res. 1982.

2. Bennetzen JL & Hall BD. Codon selection in yeast. J Biol Chem. 1982.

3. Sharp PM & Li WH. The codon adaptation index - a measure of directional synonymous codon usage bias, and its potential applications. Nucleic Acids Res. 1987.

4. Sandhu KS et al. GASCO: genetic algorithm simulation for codon optimization. In Silico Biol. 2008.

5. Blażej P et al. Optimization of the standard genetic code according to three codon positions using an evolutionary algorithm. PLoS One. 2018.

6. Sen A et al. Codon optimization: a mathematical programing approach. Bioinformatics. 2020.

7. Taneda A & Asai K. COSMO: A dynamic programming algorithm for multicriteria codon optimization. Comput Struct Biotechnol J. 2020.

8. Zhang H et al. Algorithm for optimized mRNA design improves stability and immunogenicity. Nature. 2023.

9. Kalyanmoy Deb et al. A Fast and Elitist Multiobjective Genetic Algorithm: NSGA-II. IEEE Transactions on Evolutionary Computation. 2002.

10. Kalyanmoy Deb & Himanshu Jain. An Evolutionary Many-Objective Optimization Algorithm Using Reference-point Based Non-dominated Sorting Approach. IEEE. 2013.

11. Amin Ibrahim et al. EliteNSGA-III: An Improved Evolutionary Many-Objective Optimization Algorithm. 2016.

12. Fath S et al. Multiparameter RNA and codon optimization: a standardized tool to assess and enhance autologous mammalian gene expression. PLoS One. 2011.

13. Papamichail D et al. Codon Context Optimization in Synthetic Gene Design. IEEE/ACM Trans Comput Biol Bioinform. 2018.

14. M. Gardiner-Garden & M. Frommer. CpG Islands in Vertebrate Genomes. J Mol Biol. 1987.

15. Jain R et al. ICOR: improving codon optimization with recurrent neural networks. BMC Bioinformatics. 2023.

16. Goulet DR et al. Codon Optimization Using a Recurrent Neural Network. J Comput Biol. 2023.

17. Fu H et al. Codon optimization with deep learning to enhance protein expression. Sci Rep. 2020.

18. Gong H et al. Integrated mRNA sequence optimization using deep learning. Brief Bioinform. 2023.

19. Ingolia, Nicholas T., et al. Genome-wide analysis in vivo of translation with nucleotide resolution using ribosome profiling. Science. 2009.

20. Brar, Gloria A., and Jonathan S. Weissman. Ribosome profiling reveals the what, when, where and how of protein synthesis. Nature reviews Molecular cell biology. 2015.

21. Mustroph, Angelika, et al. Isolation and analysis of mRNAs from specific cell types of plants by ribosome immunopurification. Plant Organogenesis: Methods and Protocols. 2013.

22. Chaudhury S, Lyskov S & Gray JJ. PyRosetta: a script-based interface for implementing molecular modeling algorithms using Rosetta. Bioinformatics. 2010.

23. Dauparas J et al. Robust deep learning-based protein sequence design using ProteinMPNN. Science. 2022.

24. Pédelacq JD et al. Engineering and characterization of a superfolder green fluorescent protein. Nat Biotechnol. 2006.

25. Kuhlman B, Jacobs T & Linskey T. Computational Design of Protein Linkers. Methods Mol Biol. 2016.

26. Chen X, Zaro JL & Shen WC. Fusion protein linkers: property, design and functionality. Adv Drug Deliv Rev. 2013.

27. Liu W et al. TranslatomeDB: a comprehensive database and cloud-based analysis platform for translatome sequencing data. Nucleic Acids Res. 2018.

28. Gu X, Qi Y & El-Kebir M. DERNA Enables Pareto Optimal RNA Design. J Comput Biol. 2024.

29. Mauger DM et al. mRNA structure regulates protein expression through changes in functional half-life. Proc Natl Acad Sci U S A. 2019.

30. Hargrove JL & Schmidt FH. The role of mRNA and protein stability in gene expression. FASEB J. 1989.

31. Lorenz R et al. ViennaRNA Package 2.0. Algorithms Mol Biol. 2011.

32. Sun, D. Design of time-delayed safety switches for CRISPR gene therapy. Sci Rep. 2021.

